# Bottom-up parameterization of enzyme rate constants: Reconciling inconsistent data

**DOI:** 10.1101/2023.12.05.570215

**Authors:** Daniel C. Zielinski, Marta R.A. Matos, James E. de Bree, Kevin Glass, Nikolaus Sonnenschein, Bernhard O. Palsson

## Abstract

Kinetic models of enzymes have a long history of use for studying complex metabolic systems and designing production strains. Given the availability of enzyme kinetic data from historical experiments and machine learning estimation tools, a straightforward modeling approach is to assemble kinetic data enzyme by enzyme until a desired scale is reached. However, this type of ‘bottom up’ parameterization of kinetic models has been difficult due to a number of issues including gaps in kinetic parameters, the complexity of enzyme mechanisms, inconsistencies between parameters obtained from different sources, and *in vitro-in vivo* differences. Here, we present a computational workflow for the robust estimation of kinetic parameters for detailed mass action enzyme models while taking into account parameter uncertainty. The resulting software package, termed MASSef (the Mass Action Stoichiometry Simulation Enzyme Fitting package), can handle standard ‘macroscopic’ kinetic parameters, including K_m_, k_cat_, K_i_, K_eq_, and n_h_, as well as diverse reaction mechanisms defined in terms of mass action reactions and ‘microscopic’ rate constants. We provide three enzyme case studies demonstrating that this approach can identify and reconcile inconsistent data either within *in vitro* experiments or between *in vitro* and *in vivo* enzyme function. The code and case studies are provided in the MASSef package built on top of the MASS Toolbox in Mathematica. This work builds on the legacy of knowledge on kinetic behavior of enzymes by enabling robust parameterization of enzyme kinetic models at scale utilizing the abundance of historical literature data and machine learning parameter estimates.

**Author Summary:** Detailed kinetic models of metabolism offer the promise of enabling new predictions of metabolic behavior and prospective design of metabolic function. However, parameterizing such models remains a substantial challenge. Historically, the kinetics of many enzymes have been measured using *in vitro* assays, but integrating this data into consistent large-scale models and filling gaps in available data has been a primary difficulty. Here, we provide an algorithmic approach to parameterize enzyme kinetic models using diverse enzyme kinetic data. The approach reconciles inconsistent data and addresses the issue of gaps in available data implicitly through sampling alternative parameter sets. We provide a number of case studies demonstrating the approach on different enzymes. This work empowers the use of the large amount of historical and machine learning-estimated enzyme data and will aid in the construction of biochemically-accurate models of metabolism.

## Introduction

There has been a resurgence of interest in the construction of large-scale kinetic models of metabolism for model organisms in recent years(1–3). These models hold promise in a number of applications that constraint-based models have difficulty addressing, such as understanding quantitatively how metabolite levels control metabolic flux across experimental conditions(4–7). However, the primary issue impeding the development of practical large-scale kinetic models of metabolism is the need for a large number of kinetic parameters, the vast majority of which have not been experimentally measured(8).

To address this parameterization challenge, a number of approaches have been developed(9,10). These approaches can be loosely classified into parameter sampling, “top-down” parameterization, and “bottom-up” parameterization methods. A particular model parameterization workflow may integrate multiple categories of these methods. Sampling methods randomly select parameters in particular expected ranges, sometimes with constraints such as thermodynamic consistency enforced in the sampling procedure(11–13). Top-down parameterization methods use data on the kinetic or steady-state behavior of the entire system and parameterize the entire model simultaneously to match this data(14–16). Bayesian methods have been proposed as a flexible and powerful tool for parameterization (17,18). Both sampling and top-down methods can efficiently define the large number of parameters required by a kinetic model, although this can require substantial computation. Further, a number of software tools are available for parameterization using these methods(19–22). However, performance of such methods can suffer due to sampling too wide a parameter space or having too many parameters for the limited amount of data.

Bottom-up methods on the other hand use data related to the individual components of the network to construct a model piece by piece(23,24). Various fitting methods have been developed to determine kinetic parameters that match enzyme kinetic experimental data such as initial rate and continuous experiments to user-defined enzyme mechanisms (25–27). Bottom-up methods have the advantage of utilizing the extensive amount of historical enzyme data(28,29) as well as machine learning estimates(30,31). However, there are a number of difficulties with bottom-up construction of kinetic models that have impeded their development. First, the majority of enzymes do not have detailed kinetic assays performed to measure the required kinetic parameters to construct such models. As a result, kinetic models are often highly underdetermined and methods to estimate missing parameters must be developed(32). Also, enzyme kinetics are often measured under non-physiological conditions, raising questions of their relevance to modeling *in vivo* systems(24,33–35). However, high-throughput studies have shown substantial correlations between *in vivo* and *in vitro* kinetic parameters, supporting the use of available enzyme kinetic data(36,37). Still, parameters from disparate sources may be inconsistent, requiring a framework to integrate this data(11,38). If these challenges can be met, bottom-up methods may be promising complementary alternatives for construction of large-scale kinetic models of metabolism if these challenges can be overcome.

Here, we present a computational workflow that attempts to address a number of issues with bottom-up construction of kinetic models of metabolism. This workflow parameterizes individual enzyme kinetic models to fit a variety of measured enzyme data types, attempts to reconcile inconsistent data, and accounts for parameter uncertainty. To aid this, we utilize flexible user-defined microscopic mass action reaction mechanisms that can account for the vast majority of observed kinetic behavior and can be extended to the necessary resolution on an enzyme-by-enzyme basis. This workflow is implemented as a software package in Mathematica termed MASSef (Mass Action Stoichiometric Simulation Enzyme Fitting). There are three core features of this software package: 1) a symbolic algebra system that generates equations for comparison with measured data based upon the enzyme mechanism, 2) robust nonlinear optimization to fit the model to data, and 3) a workflow that perturbs the fitting problem to characterize uncertainty in the parameters. We first present an overview of the computational workflow for parameter fitting before discussing the details of individual components, and then we present a series of case studies demonstrating the workflow for enzymes with different available kinetic data.

## Results

### Overview of enzyme kinetic parameter fitting pipeline

The parameterization workflow is divided into six steps (**Fig. 1**). 1) Gather available enzyme kinetic data and associated experimental conditions. 2) Construct a mass action enzyme mechanism, consisting of known individual reaction steps involved in catalysis and regulation of the enzyme. 3) Based upon the enzyme mechanism, define kinetic equations corresponding to the macroscopic kinetic data types (e.g. K_M_, k_cat_) to be fit. 4) Pre-process the kinetic data to allow direct comparison to the equations defined in Step 3. 5) Fit the equations defined in step 3 to the processed data in Step 4 using a nonlinear least squares optimization approach to obtain a locally optimal set of microscopic rate constants that reproduces the experimental data. Data priorities specified by the user determine the weight of each data point in the fitting procedure. This optimization problem is solved N times with pseudo-random start points to account for the under-determined nature of the problem. 6) Cluster the resulting rate constant sets to identify a reduced number of characteristic rate constant sets for the enzyme, each of which reproduces the experimentally-determined kinetic behavior of the enzyme. Finally, fit performance is summarized, effective macroscopic kinetic parameters are recalculated from the fitted microscopic rate constants, and the parameterized model can be exported as text files for modeling in desired kinetic simulation software. We discuss the details of each step below.

**Figure 1:**
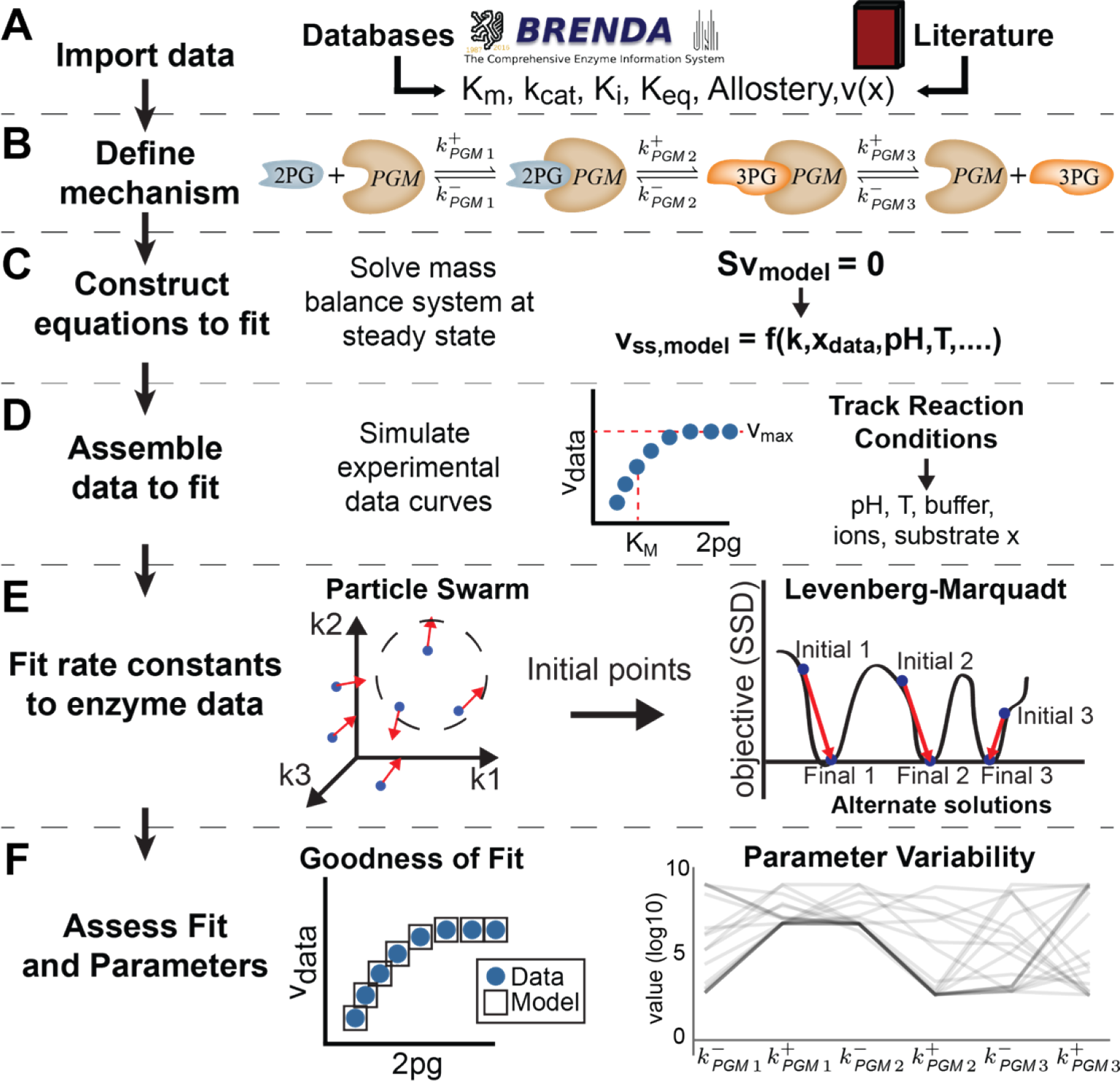
Workflow for parameterization of rate constants using enzyme kinetic data. This workflow consists of six steps: A) Gathering and curating kinetic data, B) Defining an enzyme mechanism, C) Defining equations that relate model behavior to each data type, D) Processing kinetic data and correcting it to *in vivo*-like conditions, E) A nonlinear least-squares optimization to identify rate constant sets that fit available kinetic data, F) Assessing the fit for goodness of fit and parameter uncertainty, as well as calculating clusters of similar rate constant set results.

### Step 1: Preparation of enzyme kinetic data

First, kinetic data on enzymes is gathered, curated, and placed into a table in a standard format (**Fig. 1A**). Data types currently handled include enzyme structure, reaction stoichiometry, kinetic reaction mechanism, reaction equilibrium constant (K_eq_), dissociation constants (K_d_), and standard initial rate kinetic assay constants such as the Michaelis-Menten constant, K_M_, the turnover rate, k_cat_, and inhibition constants, K_i_. Additional data types, such as *in vivo* flux data, can be utilized as well, provided that a comparison equation is specified by the user in step 3. Experimental condition data such as co-substrate concentrations, pH and temperature are also extracted for use in data adjustments to *in vivo*-like conditions, as discussed later. Weights for each data point are determined by the user. This allows the user to decide how heavily to consider data points that may be conflicting with other measurements or have reliability concerns.

### Step 2: Specify the enzyme reaction mechanism

A mechanism for the enzymatic reaction, consisting of individual reaction steps that describe the binding of the enzyme to substrates, catalysis, and product release, is then specified by the user (**Fig. 1B**). Each of these steps is modeled using mass action rate laws. Since the reaction steps do not proceed through a single transition state, these are not true elementary reactions. Instead, we refer to them using Cleland’s nomenclature(39) as microscopic reaction steps, with microscopic kinetic rate constants, contrasted with macroscopic kinetic parameters such as Michaelis constants K_M_.

The method is flexible to various reaction schemes, such as sequential versus random binding orders, ping pong mechanisms, and slow enzyme transitions. Both reversible and irreversible reactions can be specified, but fully reversible mechanisms are recommended for later use of thermodynamic Haldane relationships(40). Generally, protons and water are assumed constants and excluded from reaction mechanisms by convention. At this stage, individual catalytic tracks and thermodynamic cycles can be defined by the user to serve as thermodynamic constraints, such as Haldane constraints arising from the First Law of Thermodynamics. A catalytic track consists of a particular set of catalytic microscopic reaction steps that convert substrates into products. Haldane constraints, which consist of a multiplication of equilibrium constants along a particular catalytic track, are later fit to be equal to the overall reaction K_eq_. In addition, small molecule enzyme inhibition and activation can be modeled by adding the corresponding microscopic binding reaction step(s). For enzyme competitive, uncompetitive, and mixed inhibition mechanisms, these microscopic reactions are added automatically if the inhibitor and respective affected metabolites are specified.

We note that although the mechanism is specified in terms of mass action reactions, an overall rate equation is derived based on a quasi-steady state assumption, and either the original mass action equations or this overall rate equation can be used for downstream kinetic modeling, as the user desires.

### Step 3: Constructing symbolic comparison equations

We then set up the equations that are used to fit the kinetic data (**Fig. 1C**). These equations are functions of microscopic rate constants and in some cases of metabolite concentrations as well. For K_eq_ values, Haldane relationships are defined in terms of individual rate constants for each catalytic track through the enzyme mechanism. For k_cat_, K_M_, and K_i_ values, equations are derived from the overall steady-state rate equation of the reaction, v_ss_, as described in the methods. The equation for the steady-state flux, v_ss_, is found by solving the system of mass balance equations at steady-state along with a total enzyme sum equation representing total enzyme conservation (see **Methods** for details and **Supplementary Information** for a case study). Other data types such as dissociation constants can be used as well, if the user specifies a corresponding equation relating the data type to the enzyme microscopic rate constants.

### Step 4a: Preparing kinetic data for fitting

The macroscopic kinetic data, such as K_eq_, k_cat_, K_M_, and K_i_, are then processed into a form that enables direct comparison to model behavior (**Fig. 1D**). For reaction K_eq_ values, the processed form is simply the K_eq_ value calculated under experimental conditions (pH, IS, and T). For k_cat_, the processed form is the value along with the measured experimental conditions, including substrate concentrations. If substrate concentrations are not available, a concentration that is likely to be saturating is assumed (i.e. 1 M), in order to represent the excess concentrations typically used in the measurement of turnover rates. In these cases, it is recommended that K_M_ values be specified as well to ensure this constraint is satisfied by the final parameter set. For K_M_ and K_i_, an initial rate curve is generated using the classical Michaelis-Menten equation with the K_M_ and K_i_ value substituted when applicable. This curve is simulated at substrate concentrations an order of magnitude above and below the measured K_M_ value. Other experimental conditions such as co-substrate concentration and media conditions are reported when available. Co-substrate concentration values in particular are substituted in the equations used to fit the K_M_ and K_i_ data. This procedure attempts to simulate the original experiment. However, in lieu of this experimental plot simulation procedure, raw data could be used in principle as well.

To deal with uncertainty in macroscopic kinetic data, upper and lower bounds can be specified and data sampled from a normal distribution with a given mean and standard deviation. In the case of unknown macroscopic kinetic data, a uniform distribution spanning a characteristic range for the data type may be sampled. For example, K_m_ values tend to fall in the range of 10^−7^ to 10^−2^ M, while k_cat_ values may fall in the range of 10 to 10^5^ s^-1^. When sampling is performed, a number of samples is defined, and a data set is created for each sample point.

### Step 4b: Correcting data for in vitro *to* in vivo *differences*

To correct these values for *in vitro* to *in vivo* differences, several adjustments to the data are implemented. First, k_cat_ can be adjusted to *in vivo* temperature from *in vitro* conditions using a user defined Q10 value for the enzyme, which is specified by the user but has a default value of 2.5, typical for metabolic enzymes(41,42). Effects of temperature on K_m_ are thought to be less substantial and thus are currently ignored(43). Further, for each data type, concentrations can be corrected to the more accurate chemical activities using a Debye-Huckel model for the activity coefficient given the ionic strength under experimental conditions, though currently this must be performed by the user(44). Reaction equilibrium constants can be calculated at a specific pH and ionic strength as well using published tools(45). Finally, inhibitory effects of pH changes on enzyme behavior can be modeled by adding proton binding and dissociation reactions to the enzyme mechanism with associated dissociation constants, as has been done in the enzymology literature(46,47).

### Step 5: Two-stage randomized fitting of microscopic rate constants to macroscopic kinetic data

Once both the data and equations have been prepared, these are passed to a two-stage nonlinear least squares optimization procedure (**Fig. 1E**). The target values are given by the data, while the model values are given by the equations with the experimental conditions substituted into the equations, leaving them functions of the microscopic rate constants alone. The fitting procedure then yields microscopic rate constant values that cause the enzyme model to reproduce the measured data. As an objective, we use the absolute difference between the logarithm of the model-predicted value and the logarithm of the data, multiplied by the user-defined weights on each data value. We use bounds on the microscopic rate constants of 10^−6^ and 10^9^ s^-1^ based on typically assumed limits of diffusion and an arbitrarily slow lower bound(48). These rate constant bounds may not be relevant for reactions not involving association or dissociation. However, we have not found these bounds to affect the fitting error for the cases we have examined thus far, and the lower bound in particular is rarely hit in practice. Additionally, to address the broad scale of potential rate constants, we log transform the rate constants during this optimization. Furthermore, weights can be assigned to each data point for balancing the contribution of each data point to the fitting objective function or to reflect confidence in particular measurements.

However, the equations are highly nonlinear, causing many optimization algorithms to fail to converge to an acceptable fit. To address this challenge, we first run a non-derivative based particle swarm optimization (PSO) to find rate constant sets that fit the data well enough to serve as initial points for a more precise derivative-based optimization. The second optimization is a Levenberg-Marquadt (LMA) derivative-based optimization with the same log-parameter scaling and objective. Resulting fits are examined for total residual, and points that fit satisfactorily are kept. These rate constants are then once more substituted into the equations for the kinetic data to verify that this kinetic data used to fit the model can be reproduced.

Importantly, as PSO is a randomized algorithm, initial points passed to LMA are different each time that the algorithm is run. For typical underdetermined systems, different rate constant sets are returned by the optimization every time, each of which fits the kinetic data equivalently well. This effectively samples the rate constant space in the prevalent cases where rate constants are not uniquely specified by the available kinetic data. Thus, this procedure inherently addresses the underdetermined system issue in parameterization of enzyme reaction mechanisms.

### Step 6: Cluster parameters to extract characteristic rate constant sets

As initial points to the LMA optimization are defined pseudo-randomly based on the PSO optimization results, the resulting rate constants are generally different in each optimization. The fitting procedure is then repeated a number of times until qualitatively new alternative rate constant sets are no longer found (**Fig. 1F**). To determine convergence, clustering is used to identify whether new clusters have been formed by addition of new fitted points, or whether these points fall within existing clusters. Then, once convergence has been reached, characteristic rate constants sets for the enzyme are selected based on proximity to the centroids of these clusters. These centroids then serve as candidate rate constant sets for the enzyme, each of which reproduces measured kinetic data for the enzyme equivalently well.

After the parameters are fit, the best fit parameter set is evaluated. The macroscopic parameters corresponding to the fit microscopic rate constants are recalculated and compared to the data used in training to assess final error on each parameter. Centroid cluster parameter sets can be exported to text files for import into desired kinetic modeling packages, such as the MASSpy kinetic modeling package in Python(49).

### Case studies demonstrating the ability of MASSef to fit specific data types

Having described the workflow for parameterizing microscopic mass action reaction mechanisms using enzyme kinetic data, we now present a series of case studies demonstrating the workflow in action when different types of data are available.

### Case study 1: Demonstrating the fitting procedure on k_cat_, K_m_, and K_eq_ data for Glyceraldehyde-3-phosphate Dehydrogenase

We first demonstrate the basic capabilities of the fitting procedure, using the case of the glycolytic reaction Glyceraldehyde-3-phosphate Dehydrogenase (GAPDH) from *E. coli* (**Fig. 2**). This reaction catalyzes the substrate level phosphorylation of glyceraldehyde-3-phosphate (g3p) to produce 1,3-diphosphoglycerate (13dpg) and NADH. The enzyme has reported K_m_ values of 0.89mM, 0.045mM, and 0.53mM for glycerol-3-phosphate, NAD, and inorganic phosphate (P_i_), respectively(50) (**Fig. 2B**). Also reported in the study was a forward k_cat_ of 268 s^-1^ at 295K and a dissociation constant K_d_ for NAD of 0.00032mM. A K_eq_ value of 0.452 was extract from eQuilibrator at a pH of 7 and ionic strength of 0.25M. While the substrate binding order in *E. coli* has not been experimentally determined, it has been reported in humans to be an ordered Bi Bi mechanism with a binding order NAD, then g3p, then P_i_, and a release order of 13dpg followed by NADH(51). This mechanism was constructed and fit to the data using the MASSef workflow, and rate constants enabling the model to perfectly fit the data were found (**Fig. 2C**). Rate constants were not uniquely identified, but instead were constrained to certain ranges (**Fig. 2D**). Due to the presence of more data on the forward direction than the reverse, the microscopic rate constants associated to the substrate binding steps are constrained to a greater degree. Examination of good fitting rate constant sets showed nonlinear dependencies between the rate constants, a reflection of the alternate possible rate constant sets that can equivalently satisfy the constraints on enzyme function places by measured turnover rates and Michaelis-Menten constants (**Fig. 2E**). Clustering of the identified rate constants then extracts sets of characteristic rate constants with particular nonlinear dependencies (**Fig. 2F**).

**Figure 2:**
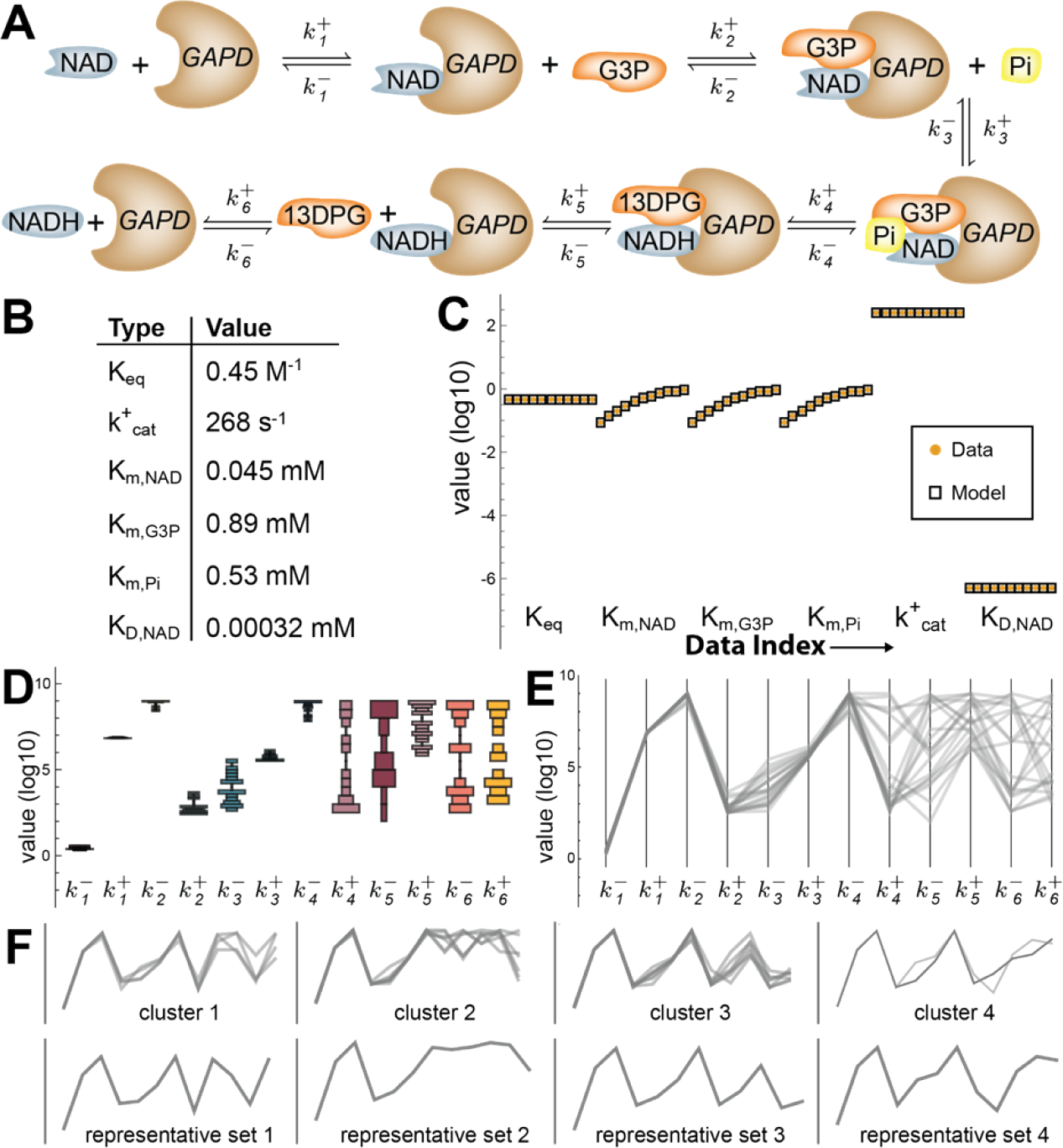
Demonstrating the fitting procedure for Glyceraldehyde-3-phosphate Dehydrogenase (GAPDH) from *E. coli*. A) Enzyme mechanism for GAPDH. B) Available kinetic data for GAPDH. C) Best fit results for GAPDH and resulting kinetic parameters calculated using symbolic equations for these parameters. Target data is shown as circles, while the model results are shown as boxes for clarity. Data is ordered by index as it is generated from the data simulating equations by type of data (e.g. K_eq_, K_m_, k_cat_). Data is replicated for certain data types (K_eq_, k_cat_, K_D_) to balance the contribution of error on these data points to the objective function with that of K_M_ data. For this enzyme, the fit between model and data was tight, as no discrepancies are apparent. D) Distribution plot of rate constant sets obtained from 100 optimizations showing parameter ranges E) Rate constant sets showing connections between rate constants F) Clustering of rate constant sets reveals nonlinear dependencies between rate constants. The four identified clusters and the rate constant sets that are nearest to the cluster means are shown.

### Case study 2: Reconciling inconsistent data for Phosphoglycerate Mutase

We then demonstrate the ability of MASSef to reconcile data that is inherently inconsistent, using Phosphoglycerate Mutase (PGM) as a case study. This enzyme catalyzes the conversion of 2-phosphoglycerate to 3-phosphoglycerate and has well specified kinetics, with both forward and reverse k_cat_ and K_M_ defined by data (**Fig. 3A,B**). The k_cat_ data was corrected to 37°C using a Q_10_ of 2.5. The equilibrium constant for the reaction was obtained from the eQuilibrator web server(52) for pH of 7.5 and IS of 0.25M. The fitting procedure reveals that the kinetic data and K_eq_ are not kinetically consistent (**Fig. 3C**). The reason for this inconsistency is apparent once the Haldane relationship is calculated using the kinetic data and compared to the reaction equilibrium constant. The K_eq_ for the reaction is 5.3, while the K_eq_ calculated from kinetic parameters is only 1.4, indicating significant inconsistency. This type of inconsistency suggests that some data may not be trustworthy. While distinguishing good data from bad data is the task of the modeler, the workflow results in rate constants that attempt to reconcile the inconsistent data as well as possible. The resulting kinetic constants are not far from the measured values. Furthermore, different weight values can be used to weigh particular data points more heavily based on user confidence. We demonstrate several possible data point weights for this enzyme, based on individually deprioritizing K_m_s, k_cat_s, or K_eq_ data (**Fig. 3C,D,E**).

**Figure 3:**
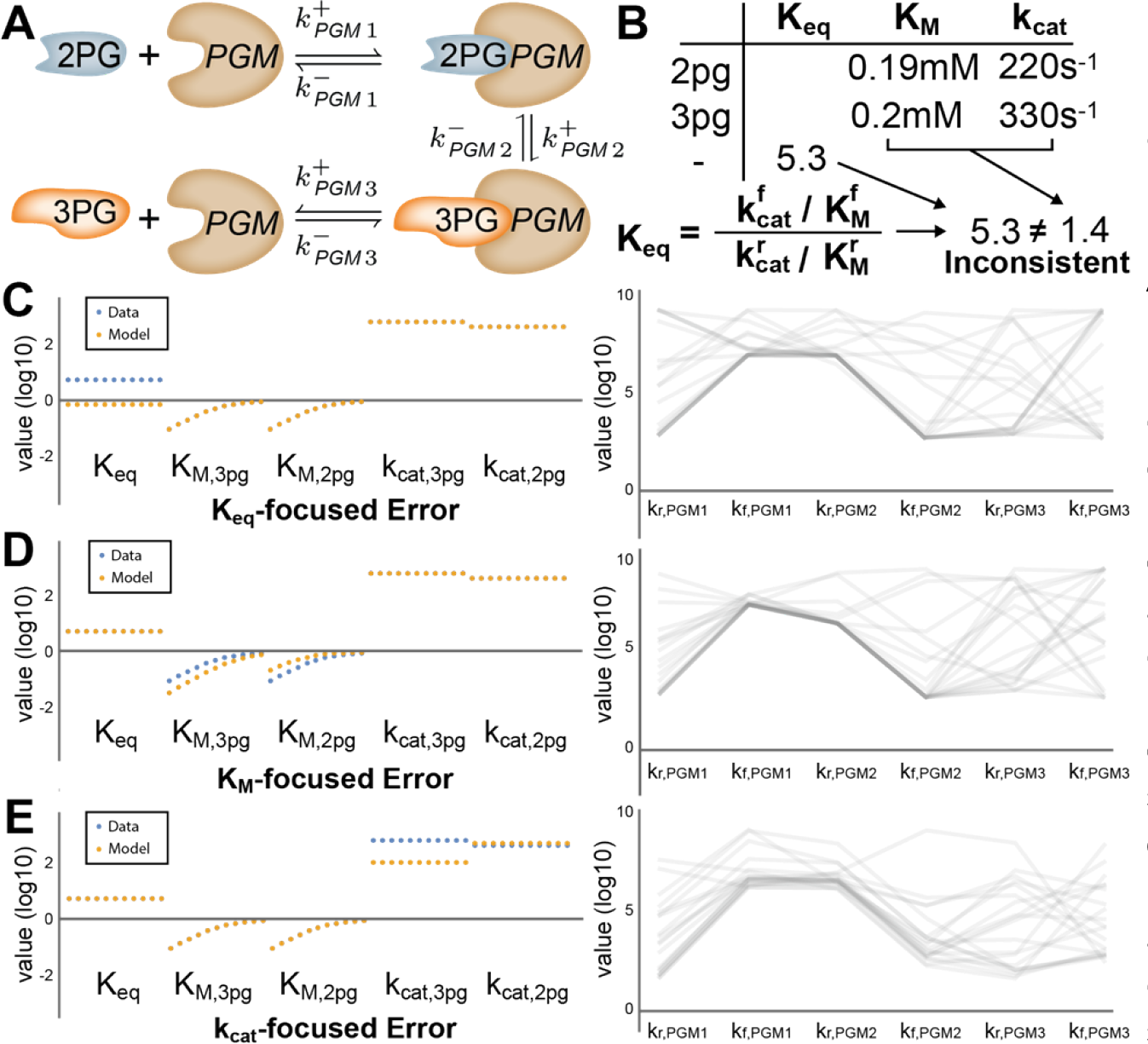
Fitting data for Phosphoglycerate Mutase (PGM) from *E. coli* shows customizable handling of inconsistent data. A) Enzyme mechanism for PGM. B) Available kinetic data for PGM. C) Best fit results for PGM and resulting kinetic parameters calculated using symbolic equations for these parameters. Error in the K_eq_ values are due to data conflicts between K_m_, k_cat_, and K_eq_ due to the Haldane constraint that relates these parameters. On the right, distribution plot of rate constant sets obtained from 100 optimizations showing parameter ranges D) Best fit results for PGM and resulting kinetic parameters calculated with data priorities scores adjusted to de-prioritize K_m_ data, which has the effect of placing the fitting error on these values. On the right, resulting rate constants show small changes as a result of the adjusted fit. E) Best fit results for PGM and resulting kinetic parameters calculated with data priorities scores adjusted to de-prioritize k_cat_ data, which has the effect of placing the fitting error on these values. On the right, resulting rate constants show small changes as a result of the adjusted fit.

### Case study 3: Incorporating in vivo flux data for Triose Phosphate Isomerase

A common issue with the use of *in vitro* data is an incompatibility with the *in vivo* requirements of pathway flux. For example, a k_cat_ may be measured inaccurately or in a non-physiological condition, resulting in less activity than required to sustain a physiological flux given a measured protein concentration. Here, we demonstrate how MASSef can incorporate *in vivo* data to obtain a set of microscopic kinetic parameters that balance *in vitro* measurements with *in vivo* requirements. We show this for the glycolytic enzyme triose-phosphate isomerase, TPI, in *E. coli*, which converts dihydroxyacetone phosphate (DHAP) to glyceraldehyde-3-phosphate (G3P) (**Fig. 4A**). The equilibrium of this reaction is slightly favored toward DHAP with a Keq of 0.11 under standard conditions, and extremely fast kinetics in the direction toward DHAP production with a catalytic efficiency near the diffusion limit, while an experimental K_M_ for DHAP was not available in the literature (**Fig. 4B**). We gathered *in vivo* data on fluxes estimated from flux balance analysis based on metabolic exchanges during aerobic growth on glucose and acetate(53), metabolite concentration data(53), and proteomics data(54) (**Fig. 4C**). We first fit the *in vitro* data alone using MASSef and found that the model was able to perfectly capture the data (**Fig. 4D**). We then added the *in vivo* data as additional target values, by asking the model to fit the *in vivo* flux given the measured metabolite and enzyme concentrations (**Fig. 4E**). The model was unable to perfectly fit both *in vitro* and *in vivo* data, indicating an inconsistency. As lower glycolysis reactions are known to be near equilibrium, which affects the thermodynamic efficiency of the enzyme(55), we hypothesized that changing the metabolite concentrations may enable a consistent fit. Indeed, by adjusting the metabolite concentrations only slightly to affect the distance of the reaction from equilibrium, by changing G3P from 0.08mM to 0.1mM on the glucose condition, we were able to find rate constants that match both *in vitro* and *in vivo* behavior (**Fig. 4F**). We recalculated the effective macroscopic parameters and found that the unadjusted *in vivo* data resulted in less efficient kinetic parameters than predicted by *in vitro* experiments, while the adjusted *in vivo* data was consistent with the *in vitro* experiments (**Fig. 4G**). The adjusted concentrations were well within apparent experimental error, especially given that condition-specific concentrations for DHAP were not available. Thus, we interpret this result to indicate that neither *in vitro* nor *in vivo* data are inherently problematic or erroneous, but rather the system is inherently highly sensitive to small changes in all parameters due to the reaction being near equilibrium where sensitivity to metabolite concentration differences is high. This work highlights the need to systematically reconcile *in vitro* and *in vivo* data to obtain consistent kinetic models, and specifically suggests the sensitivity of reactions near equilibrium to fine adjustments in metabolite concentrations as a source of potential discrepancies.

**Figure 4:**
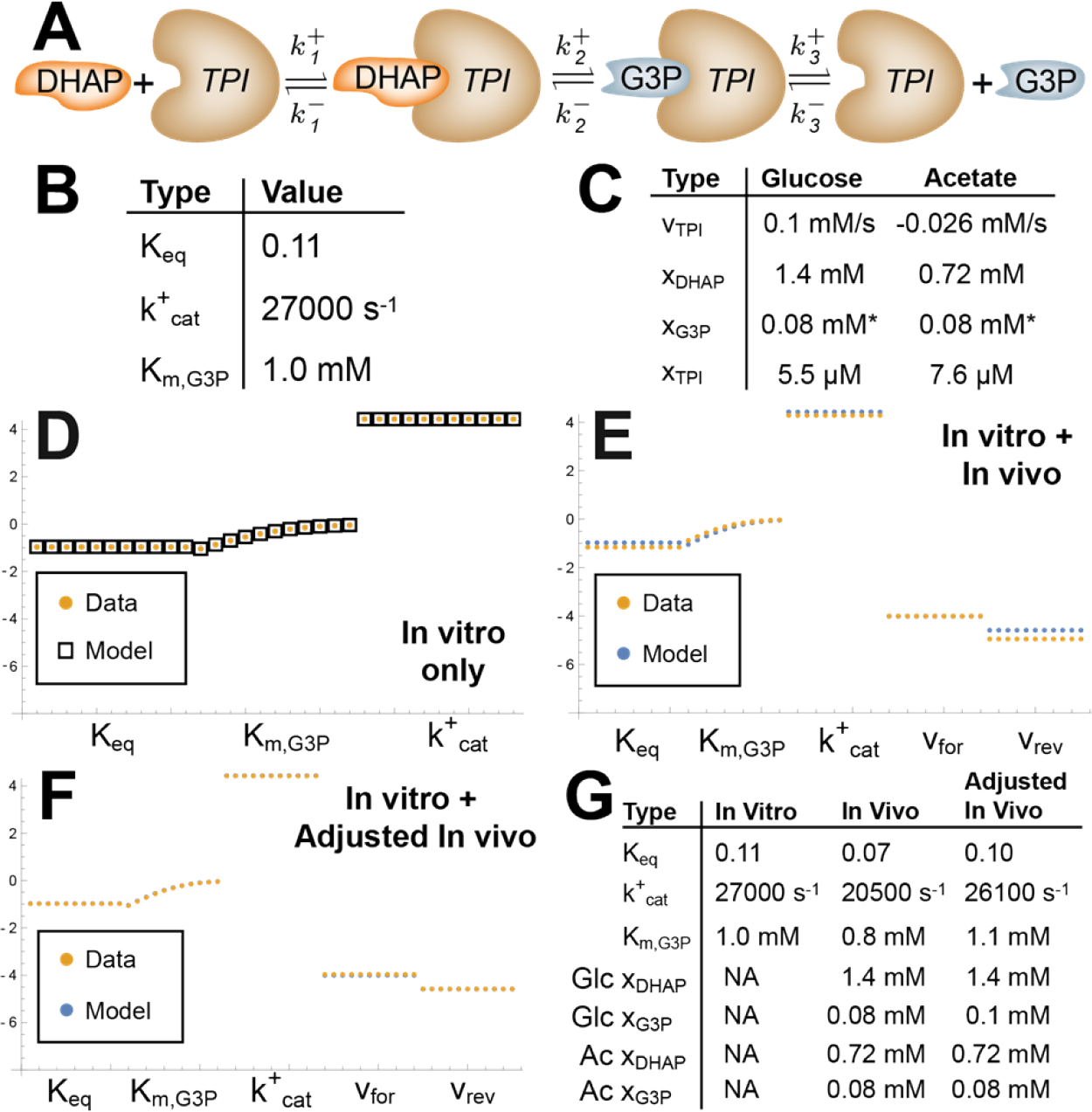
Fitting in vivo data for Triose Phosphate Isomerase (TPI) from E. coli enforces consistency of *in vitro* data with physiological function. A) Enzyme mechanism for TPI. B) Thermodynamic and In vitro kinetic data for TPI. C) *In vivo* data for TPI, including flux, metabolite concentration, and enzyme concentration data. *indicates manually adjusted value to ensure thermodynamic feasibility. D) Data fit using only thermodynamic and in vitro data. E) Data fit using thermodynamic, *in vitro*, and *in vivo* data, with *in vivo* data heavily weighted to force a tight fit. F) Data fit using thermodynamic, *in vitro*, and adjusted *in vivo* data. G) Recalculated kinetic parameters for each fit, as well as *in vivo* metabolite concentrations used in the fit.

## Discussion

In this work, we present a workflow and software tools for the parameterization of detailed kinetic models of enzyme reactions using enzyme kinetic data. This software tool has a number of powerful features. First, the workflow allows flexible user-defined reaction mechanisms and can handle the majority of common reaction schemes, including different binding orders, reaction mechanisms such as ping-pong mechanisms, inhibition schemes, protonation reactions, and allostery. Second, the workflow enables the fitting of standard kinetic data types, including thermodynamic and initial rate data, and can correct these data, to a degree, to *in vivo*-like conditions. Third, the optimization procedure can handle the inherently highly nonlinear least-squares optimization and sample rate constant sets to deal with under-determined systems.

The modeling procedure here depends on user-specified reaction mechanisms. The reaction schemes that we have tested thus far are those based on standards set by Cleland and others enzymologists. These reaction mechanisms have certain assumptions but have seen practical success in representing enzyme behavior in initial rate and progress curve experiments. There are additional levels of detail that potentially could be represented that we have not yet tested. These include additional breakdown of catalysis into individual interactions between the substrate and catalytic residues as well as breakdown of common deterministic and well-mixed assumptions due to restricted geometries, low copy numbers, channeling, stochastic behavior, proton tunneling, and other more complex kinetic phenomena. These more complex situations have been handled by others in various ways but not yet integrated into our workflow.

The current workflow has been developed to handle reaction K_eq_ values as well as enzyme initial rate data such as k_cat_, K_M_, and K_i_, due to the dominance of these data types in the enzyme kinetic literature. However, in principle this workflow could be extended to additional data types such as progress curve data and stop-flow data. The requisite for utilizing these data types is the construction of equations to be used in the least-squares optimization. For example, to compare to progress rate data, the mass balance equations for the enzyme would be integrated over time and compared to the experimental time course data. Additionally, in the current procedure we use simulated experimental curves that are effectively back-calculated for the parameters in the case of K_M_ and K_i_. This procedure was implemented because the original data curves are often not available or are inconvenient to extract from the literature. However, in principle the original data plots could be fit directly and should have mostly equivalent results to the current workflow.

Microscopic mass action enzyme systems typically have many more rate constants than kinetic data, and thus are highly underdetermined. One of the most powerful aspects of this workflow is the ability to fit rate constants for these underdetermined systems, due to the inherent parameter sampling built into the optimization procedure. This capability is enabled by the pseudo-random start points for the LMA optimization that are provided by the initial non-derivative based PSO optimization. One potential issue is to determine how completely the parameter space is being sampled. We attempt to address this by clustering the rate constants through successive optimizations, and considering the parameter space to be fully specified when new clusters are no longer found. However, this procedure cannot discount inherent bias in the optimization procedure that would lead parameters with particular biases to be oversampled. Thus, more powerful sampling methods, such as Monte Carlo sampling, would still be desirable to be implemented.

Kinetic modeling of metabolic networks has seen a resurgence in recent years, driven largely by parameterization strategies involving sampling and system-level parameter fitting. The work here is intended to increase the accessibility of a ‘bottom-up’ parameterization approach that makes use of the plethora of historical enzyme kinetic data available in the literature. As these models are parameterized enzyme-by-enzyme, this parameterization approach is expected to scale well to the network scale. Additionally, there have been recent efforts to fill gaps in critical parameters such as turnover rates at the genome-scale that should further enable these efforts(30,36). As parameterization becomes increasingly computationally feasible and biochemically accurate, practical kinetic models will likely soon become accessible and powerful tools for the systems biology community to study metabolism.

## Methods

### Setting up comparison equations

To construct comparison equations for each data type, the overall steady-state equation of the system is first solved by solving the system of equations consisting of mass balances on each species along with an expression of the total sum of enzyme:

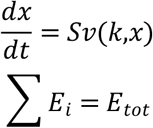

Where **S** is the reaction stoichiometry. This yields an expression for the steady state flux v_ss_ that is a function of the microscopic rate constants **k**, the metabolite concentrations **x**, and the total enzyme **E**_**tot**_. This expression can be used to derive comparison equations for specific data types as described below.

***k***_***cat***_: The enzyme turnover rate **k**_**cat**_ is a proportionality constant between the maximum rate of the enzyme **v**_**max**_ and the total enzyme concentration **E**_**tot**_. To obtain an expression that approximates **v**_**max**_, we insert concentrations of the metabolite that are assumed to be saturating, creating an approximating function **v**_**sat**_. A standard value of 1 M is used for this saturating concentration as a default, but the user can specify this value as needed. Then, the comparison equation is found by:

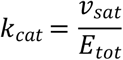

In this expression, the value of **E**_**tot**_ does not need to be specified, as **v**_**sat**_ is a function of **E**_**tot**_ as well and these terms cancel from the equation. Product and inhibitor concentrations are set to zero in these expressions, unless otherwise defined, which matches the standard conditions for initial rate experiments.

***K***_***m***_: The Michaelis constant **K**_**m**_ for a metabolite is defined as the concentration of that metabolite at which the enzyme rate is half of its maximal value.

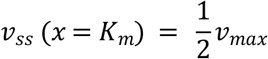

While this equation can be directly solved to generate an expression for *K*_*m*_, this can yield multiple solutions for certain reaction mechanisms. As a simplifying approach, we instead define a relative rate **v**_**rel**_

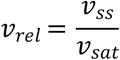

Where v_sat_ is the overall rate law with an assumed saturating concentration of the metabolites substituted in. The validity of this saturating concentration must be reassessed at the end by the user to ensure that the effective K_M_ of the enzyme is sufficiently below the saturating concentration to ensure effective saturation. In generating this expression, the concentration of any co-substrates is set to their experimental values. This expression can then be directly fit to a simulated Michaelis-Menten curve generated with the experimental **K**_**m**_ value.

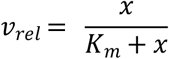

***K***_***i***_: Inhibition constants are fit analogously to **K**_**m**_ values, but additional terms are present in the simulated Michaelis-Menten curve, dependent upon the types of inhibition present. Competitive (K_ic_), uncompetitive (K_iu_), and noncompetitive schemes are currently supported.

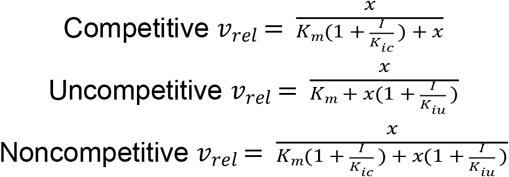

***K***_***eq***_: Reaction equilibrium constant data is fit by generating a Haldane relationship for each catalytic path through the enzyme.

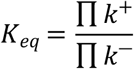

These catalytic paths are currently manually specified by the user.

### Optimization

To find microscopic rate constants that measured macroscopic enzyme kinetic data, we utilized a two-stage non-linear regression approach. Rate constants were constrained to be between 10^−6^ s^-1^ and 10^9^ s^-1^, where the upper bound is set based on the diffusion limit and the lower bound is an arbitrarily slow reaction rate. In practice, we did not observe rate constants within several orders of magnitude of the lower bound in any of the rate constant set solutions. Rate constant variables were transformed to logarithmic variables to help span the large range in possible rate constants.

Each optimization step minimizes the sum of squared residuals between the comparison equations and the data values. Priorities on each data point are specified by the user and used as a weighting on errors on those data points in the objective function. As certain data types, such as **K**_**m**_ values, consist of curves rather than single points and thus may artificially weigh more heavily in the optimization, the data points are balanced to contribute equally to the objective.

In the first optimization step, a non-derivative-based particle swarm optimization was run. Parameters for this optimization that were found to be successful across diverse data values and enzyme mechanisms are included as defaults in the fitting code, but can be changed by the user. The resulting rate constants from this initial optimization problem were then used as initial values to a second optimization problem. The second optimization was a derivative-based Levenberg-Marquadt method that refines the rate constant sets to a low sum of squared deviations to the measured data. Once again, robust options for this algorithm, such as tolerances, were identified based on performance across multiple enzyme mechanisms and are included as default options. This optimization procedure did not always return a good fit, due to the pseudo-randomness of initial points and the local optimality properties of the derivative-based optimization. Repeated runs of the fit were executed until a user-controlled number of good fits were found. Additional termination criteria around convergence of rate constants into a consistent clustering set were also tested.

The optimization procedure is implemented in Python.

### Clustering

Once a series of rate constant sets was calculated, these rate constant sets were clustered using the Mathematica FindClusters function, with the “Optimize” method, and 2 iterations. The median rate constant set for each cluster was then calculated. The rate constant set closest to the median value was selected as the characteristic rate constant set for that cluster.

### Software and Requirements

The MASSef package is available at https://github.com/opencobra/MASSef. The MASS Toolbox is available at http://opencobra.github.io/MASS-Toolbox/. The following Python packages and versions are required for the optimization:

*numpy 1.12.0*

*scipy 0.18.1*

*ecspy 1.1 (deprecated, package included internally to MASSef)*

*lmfit 0.9.5*

*Python 3.7+*

*Mathematica 10+*

## Acknowledgements

We would like to thank Zachary Haiman and Zafrin Dhali for useful discussions and testing.

## Supporting Information Captions

**Supplementary Information**. This file contains a demonstration of the enzyme fitting workflow for enolase as a case study, as well as a discussion of handling of catalytic tracks within the fitting workflow.

**Supplementary Data**. This file contains kinetic data used in the case studies as well as in vivo flux, protein and metabolite data for triose phosphate isomerase.

